# TAX1BP1 recruitment reactivates autophagy of Tau aggregates through ULK1 and TBK1

**DOI:** 10.64898/2026.02.12.705532

**Authors:** Bernd Bauer, Martina Schuschnig, Sascha Martens

**Author notes:** Correspondence should be addressed to B.B. or S.M.

## Abstract

The accumulation of protein aggregates is a hallmark of numerous neurodegenerative diseases. These aggregates frequently accumulate ubiquitin and the early autophagy factor p62 yet fail to undergo degradation. Whether such aggregates are principally susceptible to autophagic clearance is unknown. Here, we show that Tau aggregates associated with Alzheimer’s disease (AD) and other tauopathies persist due to defective autophagy initiation. This autophagy resistance correlates with a block in the recruitment of the autophagy receptor TAX1BP1 and the downstream machinery to the aggregates. Artificial targeting of TAX1BP1 to Tau aggregates restores their autophagic turnover. Mechanistically, TAX1BP1 co-activates ULK1- and TBK1-dependent signaling, thereby enabling aggregate sequestration by autophagosomes. We identify exclusion of the TAX1BP1 receptor as a breaking point at which pathological Tau aggregates evade autophagy. Furthermore, we show that Tau aggregates can be rendered susceptible to autophagy, revealing therapeutic strategies for their clearance in disease.

## Introduction

The accumulation of protein aggregates is associated with a plethora of devastating neurodegenerative diseases, such as Alzheimer’s disease (AD) and Parkinson’s disease ^1–3^. Cells possess a variety of quality control pathways that counteract protein aggregation. Among them, macroautophagy (hereafter autophagy) is seemingly particularly powerful in degrading protein aggregates because of its unique ability to sequester bulky cellular materials within double membrane vesicles, the autophagosomes. Surprisingly, it is becoming clear that solid protein aggregates including those accumulating in the cytoplasm of neurons in neurodegeneration are poor substrates for the autophagy machinery ^4–8^. This is a major conundrum in the field and is posing severe obstacles to the development of therapeutic strategies to harness autophagy for aggregate clearance.

The microtubule-binding protein Tau has been identified as the main component of neurofibrillary tangles, which are a hallmark of Alzheimer’s disease ^9^. Physiologically, Tau is a highly soluble protein regulating microtubule stability and axonal transport ^10^. Under pathological conditions, however, Tau forms first oligomeric and later amyloid fibrillar deposits in AD or other tauopathies ^9,10^. The degree of Tau fibril accumulation within the affected brain regions correlates well with the progression of the disease ^9^.

The selective autophagy pathway counteracting protein aggregation (termed aggrephagy) acts alongside the ubiquitin-proteasome system ^11,12^ in protein degradation. In aggrephagy, the oligomeric cargo receptor p62 binds the ubiquitinated cargo through its UBA domain and sequesters it, together with its interaction partner NBR1, in condensates ^13–18^. TAX1BP1 is subsequently recruited to these condensates by NBR1 and its ubiquitin binding activity. TAX1BP1 then recruits the two master regulators of selective autophagy: the ULK1 complex (ULK1c) and the kinase TBK1, to promote autophagosome biogenesis ^17,19,20^. Both, the ULK1 complex, comprised of the ULK1/2 kinase, ATG13, ATG101 and FIP200 ^21–25^ as well as TBK1 ^26–31^ can stimulate and coordinate the autophagy machinery including the lipid kinase PI3KC3-C1^32–35^, the lipidation complex ATG12-5/16L1^36^, the WIPI proteins ^37,38^ and LC3 ^39^. In line with its role as autophagy initiator by recruiting the ULK1c and TBK1 to the cargo, TAX1BP1 deficiency has been shown to result in compromised clearance of protein aggregates ^40^.

Surprisingly, many disease-associated protein aggregates, including Tau fibrils and tangles, are positive for the p62 cargo receptor and thus the early acting aggrephagy machinery, but are not efficiently cleared from cells ^41^. Using AD brain-derived Tau fibrils for reconstitutions, we have recently discovered that *in vitro* TAX1BP1 is excluded from these aggregates ^4^. This exclusion may be at least in part due to the masking of the ubiquitin marks on the Tau fibrils by the polymeric p62. Consistent with the exclusion of TAX1BP1 as an important contributor to the accumulation of Tau aggregates in AD, Tau tangles did not stain positive for TAX1BP1 in histology ^4^.

Here, we elucidate how Tau aggregates evade autophagic clearance in cells and how aggrephagy can be reactivated. We discovered that TAX1BP1 fails to target Tau aggregates in cells and that its chemically induced tethering to the aggregates restores their autophagic delivery into lysosomes by autophagy. The combined activities of the upstream autophagy factors ULK1 and TBK1 are required for the activation of Tau aggregate autophagy upon TAX1BP1 recruitment. Our study thus identifies reactivation of TAX1BP1 recruitment to protein aggregates as a promising avenue for drug development for neurodegenerative diseases.

## Results

### Tau aggregates are not an autophagy substrate in cells

To understand why protein aggregates evade autophagic degradation, we generated a cell line stably expressing an aggregation-prone variant of Tau. In line with previous studies ^7,42^, we observed the aggregation of Tau in cells upon addition of exogenous Tau fibrils, allowing us to investigate how these aggregates engage with the autophagy machinery (Figures 1A and 1B). These Tau aggregates colocalized with ubiquitin and the autophagy receptor p62, thus reproducing hallmarks of Tau aggregates in AD patients (Figure 1C) ^41,43,44^.

**Figure 1:**
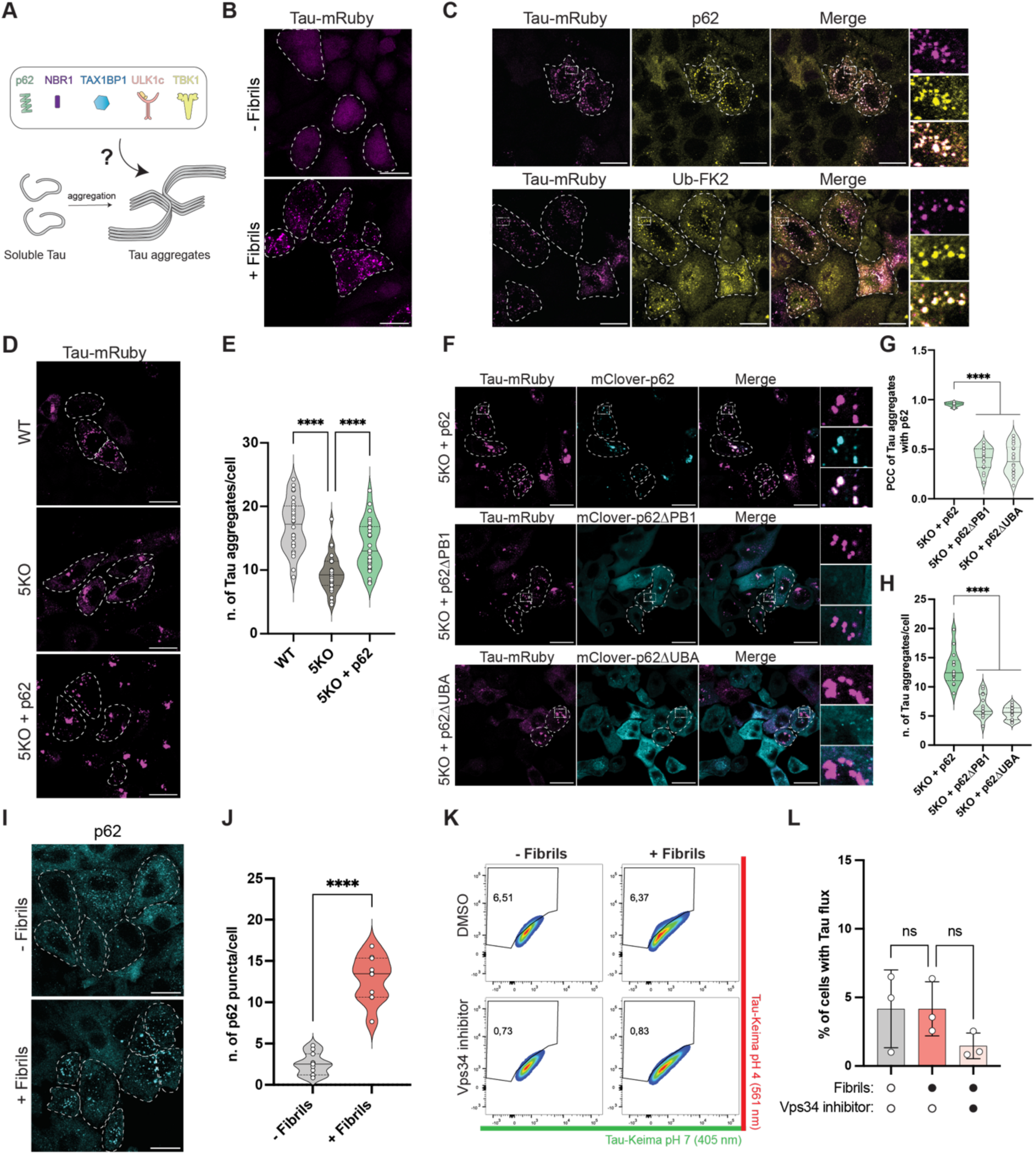
Tau aggregates are decorated by p62 but not degraded through autophagy. **A)** Schematic overview of the Tau aggregation system. **B)** Confocal images of Tau-mRuby expressing cells treated with and without *in vitro* formed Tau fibrils (scale bar = 20 µm). **C)** Immunofluorescence images of Tau, p62 and ubiquitin in cells upon treatment with Tau fibrils (scale bar = 20 µm). **D)** Confocal images of Tau in WT, 5KO and 5KO cell lines re-expressing p62 (scale bar = 20 µm). **E)** Quantification of number of Tau aggregates per cell in D. Data are presented as violin plots showing the distribution of individual data points. The solid line indicates the median; dashed lines represent the interquartile range. Each data point corresponds to the average number of aggregates of a single image. One-way ANOVA with Dunnett’s multiple comparisons test. **F)** Confocal images of 5KO Tau-mRuby cells re-expressing mClover-p62, mClover-p62ΔPB1 or mClover-p62ΔUBA (scale bar = 20 µm). **G)** Pearson’s correlation (PCC) between Tau aggregates and mClover-p62, mClover-p62ΔPB1 and mClover-p62ΔUBA in F. The data are presented as violin plots showing the distribution of individual data points. The solid line indicates the median; dashed lines represent the interquartile range. Each data point corresponds to the measured correlation within an individual cell. One-way ANOVA with Dunnett’s multiple comparisons test. **H)** Quantification of the number of Tau aggregates per cell in F. Data are presented as violin plots showing the distribution of individual data points. The solid line indicates the median; dashed lines represent the interquartile range. Each data point corresponds to the average number of aggregates of a single image. One-way ANOVA with Dunnett’s multiple comparisons test. **I)** Immunofluorescence images of p62 in cells stably expressing Tau-mRuby untreated or treated with Tau fibrils to induce Tau aggregation in the cells (scale bar = 20 µm). **J)** Quantification of the number of p62 puncta per image taken in I. Data are presented as violin plots showing the distribution of individual data points. The solid line indicates the median; dashed lines represent the interquartile range. Each data point corresponds to the average number of aggregates in a single image. Unpaired t-test. **K)** Representative fluorescence-activated cell sorting (FACS) plots of Tau flux measurements upon aggregation. **L)** Quantification of Tau flux upon Tau aggregation in C. Data are presented as mean +/- s.d. (n = 3 biologically independent experiments). One-way ANOVA with Dunnett’s multiple comparisons test. *P < 0.05, ****P < 0.0001. ns, not significant.

Next, we set out to determine how the autophagy machinery interacts with these aggregates and at which step it is stuck. First, we dissected the contribution of the main five ubiquitin-binding cargo receptors to aggregate formation by expressing Tau in cells lacking p62, NBR1, TAX1BP1, OPTN, and NDP52 (5KO) ^27^. Surprisingly, these cells had a lower number of Tau aggregates compared to wild-type (WT) cells, suggesting that the cargo receptors contribute directly to the aggregation properties of Tau (Figures 1D and 1E). The re-introduction of p62 was sufficient to increase the number of Tau aggregates in the 5KO cell line, in agreement with its role in cargo sequestration (Figures 1D and 1E) ^14,16^. p62 required both its ubiquitin-binding UBA and oligomerizing PB1 domains to colocalize with and promote Tau aggregation (Figures 1F, 1G and 1H) ^45,46^.

Since p62 marks its substrates for autophagic degradation and the aggregation of Tau led to a significant increase in the number of p62 puncta (Figures 1I and 1J), we investigated whether the increase in putative cargo coincides with the initiation of autophagic flux of Tau. To follow aggrephagy, we first established a cellular system allowing us to measure autophagic flux of protein aggregates. To this end, we generated cell lines that express Tau fused to the Keima reporter ^47^. Since the Keima reporter changes its excitation maximum based on the surrounding pH, it allows to directly measure the flux of Tau into lysosomes by flow cytometry^47^. Analogous to Tau aggregates persisting in pathologies including Tau, we did not detect any flux of Tau into lysosomes upon aggregate formation (Figure 1K and 1L).

We concluded that our cellular model reproduces key features of the aggregates found in pathologies and thus allows us to study their evasion mechanism.

### Autophagy machinery fails to assemble on Tau aggregates

Next, we sought to understand at which stage downstream of ubiquitin and p62 the autophagy cascade is blocked. To this end, we performed pelleting assays to identify which factors co-pellet with the Tau aggregates (Figure 2A). In addition to p62, the cargo receptor NBR1 also co-pelleted with the aggregates (Figures 2B and 2C). NBR1 is known to bind to p62 directly and promote its clustering by providing a high-affinity ubiquitin-binding domain ^14,15,17^. However, no additional component of the autophagy machinery co-pelleted significantly with Tau (Figures 2B and 2C).

**Figure 2:**
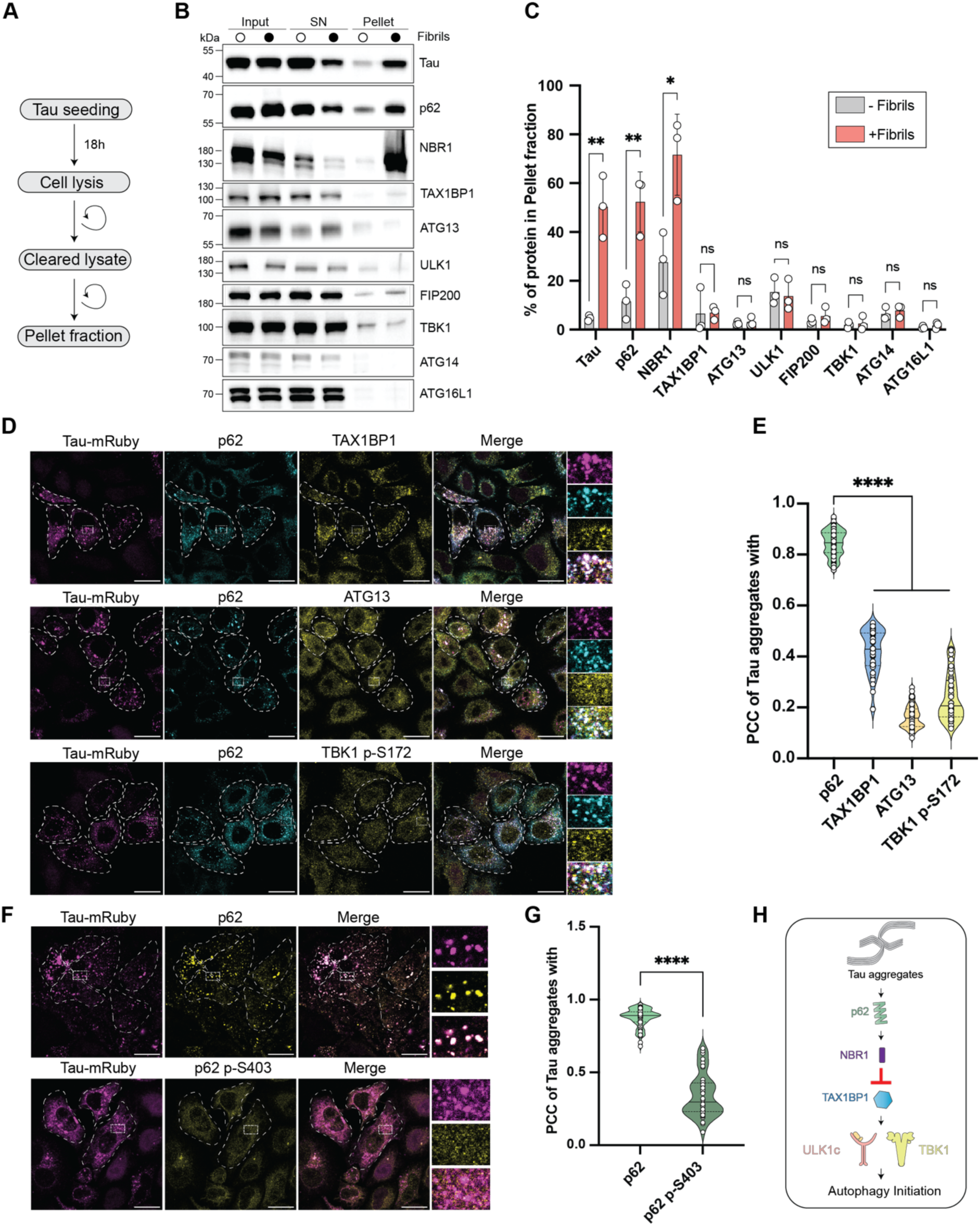
The autophagy machinery is excluded from Tau aggregates. **A)** Schematic overview of the pelleting assay. **B)** Immunoblotting for autophagy factors after Tau aggregation and the pelleting assay. **C)** Quantification of the signal in the pellet fraction in comparison to the input samples. Data are presented as mean +/- s.d. (n = 3 biologically independent experiments). Multiple unpaired t-tests. **D)** Immunofluorescence images of Tau, p62, TAX1BP1, ATG13 and TBK1 p-S172 (scale bar = 20 µm). **E)** Pearson’s correlation (PCC) between Tau aggregates and p62, TAX1BP1, ATG13 and TBK1 p-S172. Data are presented as violin plots showing the distribution of individual data points. The solid line indicates the median; dashed lines represent the interquartile range. Each data point corresponds to the measured correlation within an individual cell. One-way ANOVA with Dunnett’s multiple comparisons test. **F)** Immunofluorescence images for p62 and p62 p-S403 in cells expressing Tau-mRuby after Tau seeding (scale bar = 20 µm). **G)** Pearson’s correlation (PCC) between Tau aggregates and p62 or p62 p-S403 in I. Data are presented as violin plots showing the distribution of individual data points. The solid line indicates the median; dashed lines represent the interquartile range. Each data point corresponds to measured correlation within an individual cell. **H)** Schematic summary of the results. *P < 0.05, **P < 0.005, ****P < 0.0001. ns, not significant.

Since pelleting assays rely on cell lysis and therefore favor strong, stable interactions, we complemented them with immunofluorescence (IF) for upstream autophagy factors, including the kinase TBK1 and the ULK1-complex subunit ATG13. Activated TBK1 and ATG13 did not colocalize with the Tau aggregates (Figures 2D and 2E). Moreover, the cargo receptor TAX1BP1 showed only a minimal degree of colocalization with the Tau aggregates, consistent with the fractionation results above and our previous purely *in vitro* work showing the exclusion of TAX1BP1 from patient-derived Tau fibrils (Figures 2D and 2E)^4^. In line with a failure to recruit autophagy initiating factors, p62 associated with the Tau aggregates remained largely unphosphorylated, thus lacking an important activation mark (Figures 2F and 2G) ^16,30,48^.

Our data indicate that Tau aggregates, while recognized by p62 and NBR1, evade autophagic degradation, possibly due to the failure of the rest of the autophagy machinery to assemble on the aggregates (Figure 2H).

### Artificial tethering of TAX1BP1 initiates autophagic flux of Tau aggregates

Because the autophagy machinery fails to assemble at the Tau aggregates downstream of p62 and NBR1, we next examined whether overcoming this blockade would be sufficient to initiate autophagic flux of the aggregates. We have previously reported that the recruitment of the cargo receptor TAX1BP1 initiates autophagic degradation of p62 condensates ^19^. Since TAX1BP1 is largely excluded from Tau aggregates, we generated a cell line expressing Tau-Keima-FRB and TAX1BP1-GFP-FKBP to enforce its recruitment to the aggregates (Figure 3A).

**Figure 3:**
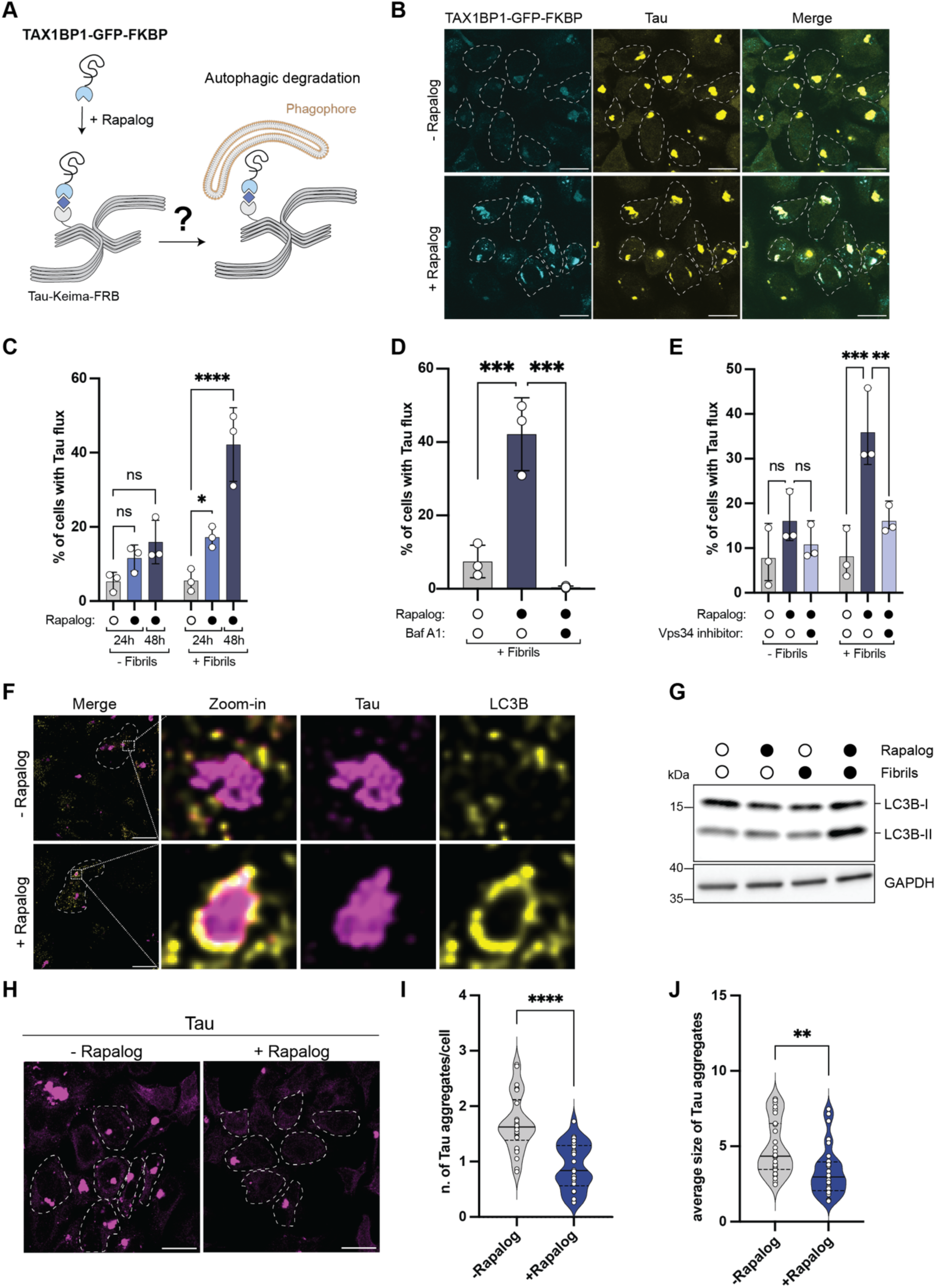
TAX1BP1 tethering initiates autophagic Tau aggregate degradation. **A)** Schematic overview of TAX1BP1 tethering to Tau aggregates through the FKBP-FRB-rapalog system. **B)** Immunofluorescence images for Tau and FKBP-GFP-TAX1BP1 untreated and treated with rapalog (scale bar = 20 µm). **C)** FACS-based measurements of Tau flux upon addition of rapalog for 24h and 48 hours with and without induction of Tau aggregation. Data are presented as mean +/- s.d. (n = 3 biologically independent experiments). Two-way ANOVA with Dunnett’s multiple comparisons test. **D)** FACS-based measurement of Tau flux upon addition of rapalog for 48 hours with and without treatment with Bafilomycin. Data are presented as mean +/- s.d. (n = 3 biologically independent experiments). One-way ANOVA with Dunnett’s multiple comparisons test. **E)** FACS-based measurement of Tau flux upon addition of rapalog for 48 hours with and without addition of the VPS34 inhibitor VPS34 IN1. Data are presented as mean +/- s.d. (n = 3 biologically independent experiments). Two-way ANOVA with Dunnett’s multiple comparisons test. **F)** Immunofluorescence images of Tau and LC3B. Images were deconvolved (scale bar = 20 µm). **G)** Immunoblotting for LC3B, showing increase levels of lipidated LC3B-II upon Tau aggregation and rapalog treatment. **H)** Immunofluorescence images for Tau following rapalog treatment (scale bar = 20 µm). **I)** Quantification of the number of Tau aggregates per cell following rapalog treatment in H. Images were taken at random positions. Data are presented as violin plots showing the distribution of individual data points. The solid line indicates the median; dashed lines represent the interquartile range. Each data point corresponds to the average number of aggregates of a single image. Unpaired t-test. **J)** Quantification of the average size of Tau aggregates per image taken in H. Images were taken at random positions. Data are presented as violin plots showing the distribution of individual data points. The solid line indicates the median; dashed lines represent the interquartile range. Each data point corresponds to the average number of aggregates of a single image. Unpaired t-test. *P < 0.05, **P < 0.005, ***P < 0.001, ****P < 0.0001. ns, not significant.

Following rapalog treatment, we observed robust colocalization between TAX1BP1 and the Tau aggregates (Figure 3B). To address changes in the flux of Tau upon TAX1BP1 tethering, we measured the ratiometric shift of the Keima probe by flow cytometry (FACS). In the presence of Tau aggregates, TAX1BP1 tethering caused a significant increase of Tau flux into lysosomes at 24 hours and particularly 48 hours following rapalog treatment (Figures 3C and 3D). Consistent with lysosomal delivery, we also detected Tau within lysosomes by IF (Figure S1). The TAX1BP1-induced increase in autophagic flux depends on autophagosome biogenesis, because its inhibition with a PI3K-complex-specific inhibitor abolished Tau flux (Figure 3E). Moreover, we frequently observed the formation of LC3B-positive, ring-shaped structures surrounding Tau aggregates in immunofluorescence, indicative of the formation of autophagosomes around these aggregates (Figure 3F). We also observed a marked increase in LC3 lipidation upon TAX1BP1 tethering by western blotting (Figure 3G). Importantly, we observed a consistent reduction in the number and size of Tau aggregates in the cells following rapalog treatment (Figures 3H, 3I and 3J).

To test whether the turnover of Tau upon artificial tethering of TAX1BP1 relies on the presence of other cargo receptors, we performed the tethering experiment in the 5KO cells. Similar to WT cells, we also observed robust Tau flux in 5KO cells, showing that TAX1BP1 does not require p62, NBR1, OPTN and NDP52 to initiate the selective autophagy of Tau aggregates (Figures S2A and S2B).

In summary, TAX1BP1 is excluded from Tau aggregates, and its recruitment is sufficient to trigger their autophagic flux into lysosomes.

### TAX1BP1-mediated aggrephagy requires both the ULK1 complex and TBK1

To investigate the mechanism of TAX1BP1-mediated aggrephagy, we first set out to identify the domains required for this activity. TAX1BP1 harbors a N-terminal SKICH domain, a C-terminal Zinc finger domain and a central coiled-coil region (Figure 4A). By re-expressing truncations of TAX1BP1 in the 5KO cell line, we found that the SKICH domain of TAX1BP1 is important for initiating Tau flux (Figures 4B and 4C).

**Figure 4:**
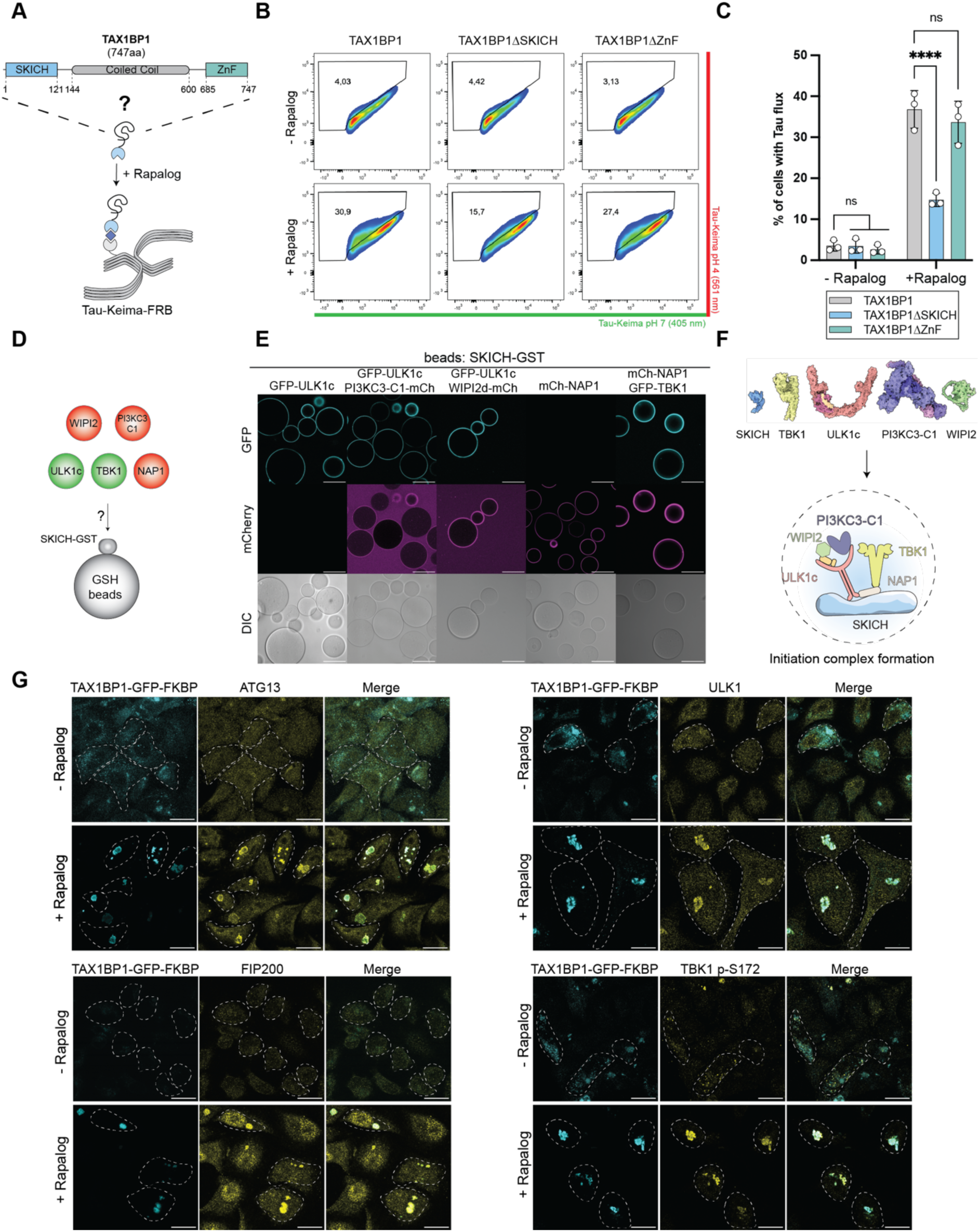
TAX1BP1 recruits the upstream autophagy machinery to Tau aggregates. **A)** Schematic overview of the domain structure of TAX1BP1 and the tethering of TAX1BP1 truncations. **B)** Representative fluorescence-activated cell sorting (FACS) plots of Tau flux measurements in cells expressing full-length TAX1BP1, TAX1BP1ΔSKICH or TAX1BP1ΔZnF. **C)** FACS-based measurement of Tau flux upon its aggregation and tethering of TAX1BP1 WT, TAX1BP1ΔSKICH and TAX1BP1ΔZnF. Data are presented as mean +/- s.d. (n = 3 biologically independent experiments). Two-way ANOVA with Dunnett’s multiple comparisons test. **D)** Schematic overview of the microscopy-based protein-protein interaction assay in E. **E)** Representative images of microscopy-based protein-protein interaction assays between the SKICH domain of TAX1BP1 on the beads and GFP-ULK1 complex, mCh-PI3K complex, mCh-WIPI2, GFP-TBK1 and mCh-NAP1 (scale bar = 80 µm). **F)** Schematic depiction of the formation of the initiation hub on the surface of the SKICH domain of TAX1BP1 together with a size comparison between the SKICH domain and the proteins involved in the formation of the complex. **G)** Immunofluorescence images of TAX1BP1, ATG13, ULK1, FIP200 and TBK1 p-S172 with and without addition of rapalog (scale bar = 20 µm). ****P < 0.0001. ns, not significant.

In line with previous studies, the SKICH domain of TAX1BP1 binds directly to the ULK1 complex through its interaction with FIP200 ^17,20,49^ and to the kinase TBK1 through its adapter proteins such as NAP1 ^19,50,51^ in a microscopy-based protein-protein interaction assay (Figures 4D and 4E). The ULK1 complex recruits further autophagy factors to the SKICH domain, including WIPI2, via its interaction with ATG13 ^52^ and the PI3K complex (PI3KC3-C1) ^29^ (Figure 4D and 4E). This also allows the formation of the previously described ULK1c:PI3KC3-C1 supercomplex^53^. The ability of the SKICH domain to recruit all major components of the upstream autophagy machinery to form an initiation complex (Figures 4F), underscores its role as a central hub for autophagy initiation. Moreover, it grants TAX1BP1 access to the two major initiation routes in selective autophagy for soluble cargo receptors: activation through the ULK1 complex and TBK1. In line with this, we observed recruitment of the entire ULK1 complex and TBK1 to the Tau aggregates upon tethering of TAX1BP1 (Figures 4G). Notably, both kinase axes are activated through recruitment by TAX1BP1, since the phosphorylation levels of TBK1 and p62 increased significantly (Figures S3A and S3B). While ULK1 itself did not exhibit changes in its phosphorylation status, ATG13, a ULK1 substrate, showed a marked increase in its phosphorylation (Figures S3A and S3B).

Next, we sought to investigate whether both kinase complexes are essential for TAX1BP1-mediated aggrephagy of Tau (Figure 5A). To this end, we analyzed the Tau flux in FIP200 and TBK1 deficient cell lines (Figure 5B). We found that, both, the ULK1 complex subunit FIP200 and TBK1 are crucial for TAX1BP1-mediated aggrephagy initiation (Figures 5C and 5D). We also tested if the direct interaction between TAX1BP1 and the upstream autophagy machinery is important for aggrephagy initiation. To this end, based on previous studies ^49,54^ we generated a SKICH domain mutant that specifically abrogates FIP200 binding, while maintaining binding to other upstream factors such as the TBK1 adapter NAP1 (Figures 5E and 5F). When tested in cells, this mutant showed a significant drop in Tau flux, suggesting that the direct interaction between the ULK1 complex subunit FIP200 and TAX1BP1 is required for aggrephagy initiation (Figure 5G). We also attempted to generate SKICH domain mutants that specifically abrogate NAP1 binding and thus TBK1 recruitment. However, all tested NAP1 binding mutants also lost their binding to FIP200, suggesting that the TAX1BP1 SKICH – FIP200 interaction modality might be more complex than generally assumed.

**Figure 5:**
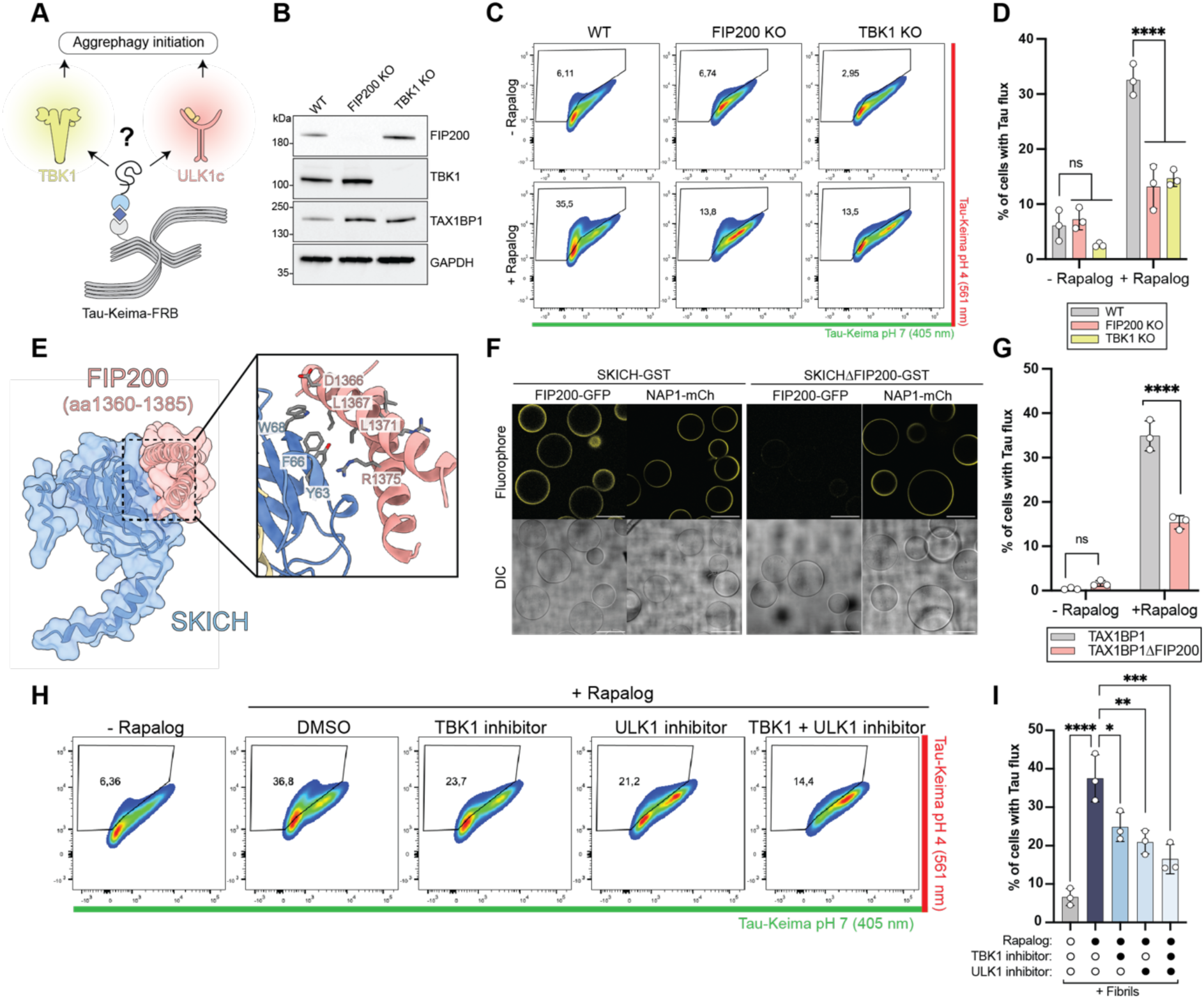
TAX1BP1-mediated aggrephagy requires the ULK1 complex and TBK1. **A)** Schematic overview of the two potential initiation axes of TAX1BP1-mediated aggrephagy. **B)** Immunoblotting for TBK1, FIP200 and TAX1BP1 to validate the TBK1 KO and FIP200 KO cell lines. **C)** Representative fluorescence-activated cell sorting (FACS) plots of Tau flux measurements in WT, FIP200 KO or TBK1 KO cells. **D)** FACS-based measurement of Tau flux following rapalog treatment in WT, FIP200 KO and TBK1 KO cell lines. Data are presented as mean +/- s.d. (n = 3 biologically independent experiments). Two-way ANOVA with Dunnett’s multiple comparisons test. **E)** AF3 prediction of the SKICH domain complex with FIP200 (aa1360-1385), including a zoom-in to better visualize the interaction interface. **F)** Representative images of microscopy-based protein-protein interaction assay comparing the binding of FIP200 and NAP1 to the WT and mutant SKICH domains (scale bar = 80 µm). **G)** FACS-based measurement of Tau flux upon aggregation and tethering of TAX1BP1 WT and TAX1BP1ΔFIP200. Data are presented as mean +/- s.d. (n = 3 biologically independent experiments). Two-way ANOVA with Dunnett’s multiple comparisons test. **H)** Representative fluorescence-activated cell sorting (FACS) plots of Tau flux measurements in cells after treatment with TBK1 and ULK1 inhibitor. **I)** FACS-based measurement of Tau flux upon aggregation and rapalog treatment without and with the addition of ULK1 and TBK1 inhibitors. Data are presented as mean +/- s.d. (n of = 3 biologically independent experiments). One-way ANOVA with Dunnett’s multiple comparisons test. *P < 0.05, **P < 0.005, ***P < 0.001, ****P < 0.0001. ns, not significant.

To further dissect the significance of the individual kinase activities of ULK1 and TBK1, we treated cells with corresponding kinase inhibitors. In line with the results obtained from the knock-out cell lines, the activities of both kinases were necessary for full TAX1BP1 mediated Tau flux (Figures 5H and 5I). Interestingly, both kinases appear to largely act in a non-redundant manner because treatment with either inhibitor led to a strong reduction of Tau flux (Figures 5H and 5I). This is noteworthy since for NDP52, the counterpart to TAX1BP1 in mitophagy, the two kinases act redundantly ^29^.

We conclude that activation of TAX1BP1-mediated autophagy of Tau aggregates requires the activities of the ULK1 complex and TBK1.

### Individually, the ULK1 complex and TBK1 are insufficient to trigger Tau flux

Next, we asked if TAX1BP1 can be bypassed by direct recruitment of the ULK1 complex or TBK1. To tether the ULK1 complex to the Tau aggregates we generated a FKBP-GFP-ATG13 expressing cell line (Figure 6A) in which upon rapalog treatment, ATG13 was recruited to the Tau aggregates (Figure 6B). In addition, the FIP200 and ULK1 subunits of the complex were co-recruited to the aggregates, as determined by cell fractionation (Figures 6C and 6D). By contrast, TBK1 was not recruited upon ATG13 tethering (Figures 6C and 6D). In agreement with the pelleting assay, we also observed the recruitment of ULK1 and FIP200 to Tau aggregates by IF (Figure 6E). Despite the assembly of the entire ULK1 complex at the site of the aggregates, we did not measure any flux of Tau by flow cytometry (Figure 6H).

**Figure 6:**
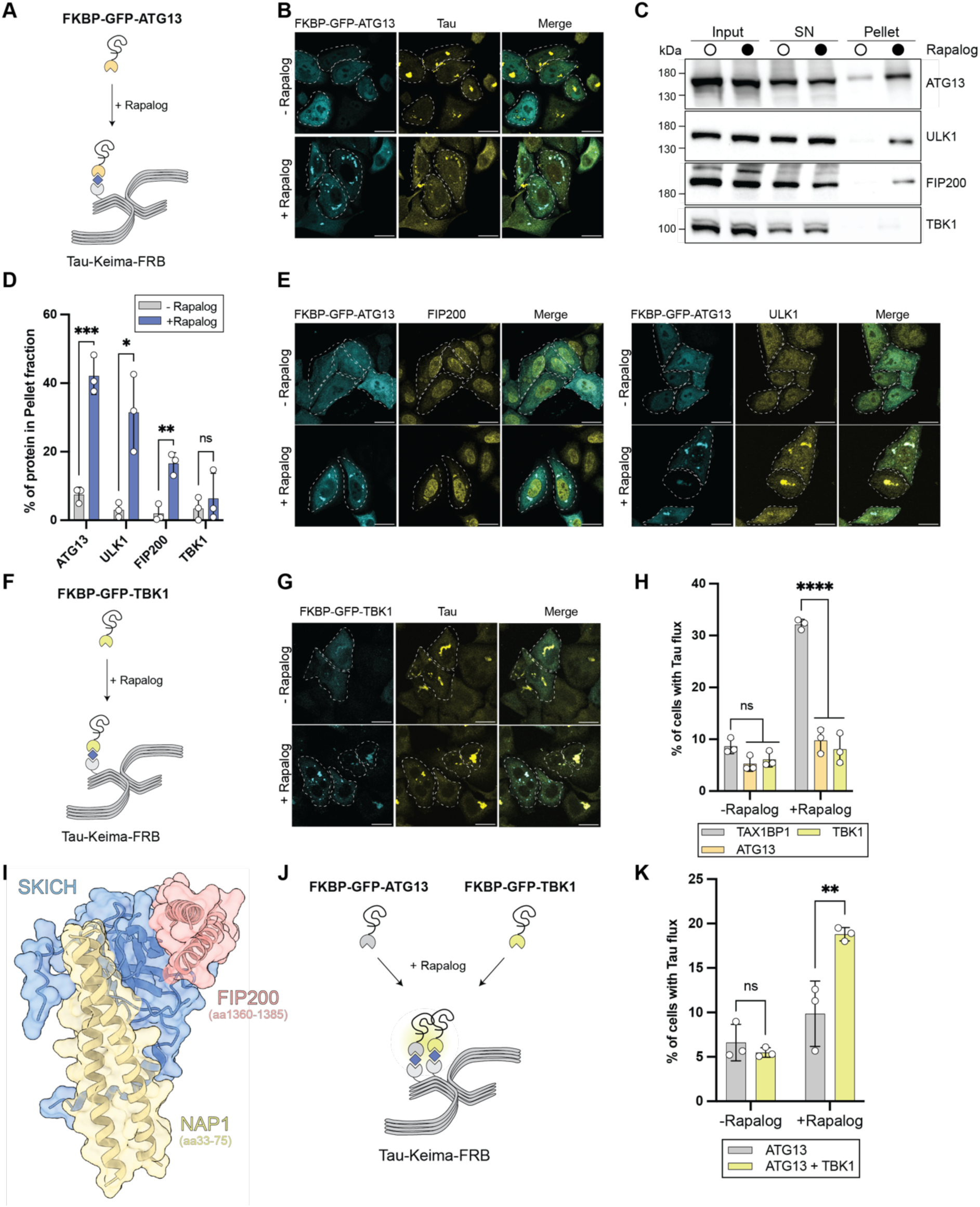
Individual tethering of the ULK1 complex or TBK1 to Tau aggregates is not sufficient to induce flux. **A)** Schematic overview of tethering ATG13 to Tau aggregates using the FKBP-FRB-Rapalog system. **B)** Immunofluorescence images of Tau and ATG13 (scale bar = 20 µm). **C)** Immunoblotting of ATG13, ULK1, FIP200 and TBK1 with and without addition of rapalog. **D)** Quantification of the fraction of protein pelleted upon rapalog treatment in C. **E)** Immunofluorescence images of Tau and ULK1 or FIP200 with and without rapalog treatment (scale bar = 20 µm) **F)** Schematic overview of tethering TBK1 to Tau aggregates using the FKBP-FRB-Rapalog system. **G)** Immunofluorescence images of Tau and TBK1 with and without rapalog treatment (scale bar = 20 µm). **H)** Comparison of the Tau flux upon tethering TAX1BP1, ATG13 or TBK1 to Tau aggregates following rapalog treatment. Data are presented as mean +/- s.d. (n = 3 biologically independent experiments). Two-way ANOVA with Dunnett’s multiple comparisons test. **I)** AF3 prediction of the ternary complex formed by the SKICH domain of TAX1BP1, aa1360-1385 of FIP200 and aa33-75 of NAP1. **J)** Schematic overview of the ATG13 and TBK1 co-tethering experiment. **K)** Comparison of the Tau flux upon tethering of ATG13 only or ATG13 and TBK1 co-tethering to the Tau aggregates. Data are presented as mean +/- s.d. (n = 3 biologically independent experiments). Two-way ANOVA with Dunnett’s multiple comparisons test. *P < 0.05, **P < 0.005, ***P < 0.001, ****P < 0.0001. ns, not significant.

Next, we tethered TBK1 to the aggregates (Figure 6F) and determined its recruitment to the Tau aggregates upon rapalog treatment using IF (Figure 6G). Similar to the results obtained by ATG13 tethering, we did not observe induction of Tau flux upon TBK1 tethering, suggesting that both kinases must be recruited and activated for efficient degradation of Tau aggregates (Figure 6H).

### ULK1c and TBK1 co-clustering promotes aggrephagy initiation

Our findings above raised the question of how the tethering of TAX1BP1 differs from tethering TBK1 or ATG13 alone. We speculated that the simultaneous recruitment of the ULK1 complex and TBK1 to the cargo by the SKICH domain of TAX1BP1 may be essential for the initiation of aggrephagy (Figure 6I). To test this hypothesis, we generated a cell line in which we could simultaneously tether the ULK1 complex and TBK1 to the Tau aggregates (Figure 6J). We observed a robust increase in Tau flux when TBK1 and ATG13 are co-tethered in contrast to ATG13 alone following rapalog treatment (Figure 6K).

In summary, we demonstrate that TAX1BP1 is sufficient to initiate autophagic degradation of Tau aggregates through local co-clustering of TBK1 and the ULK1 complex (Figure 7).

**Figure 7:**
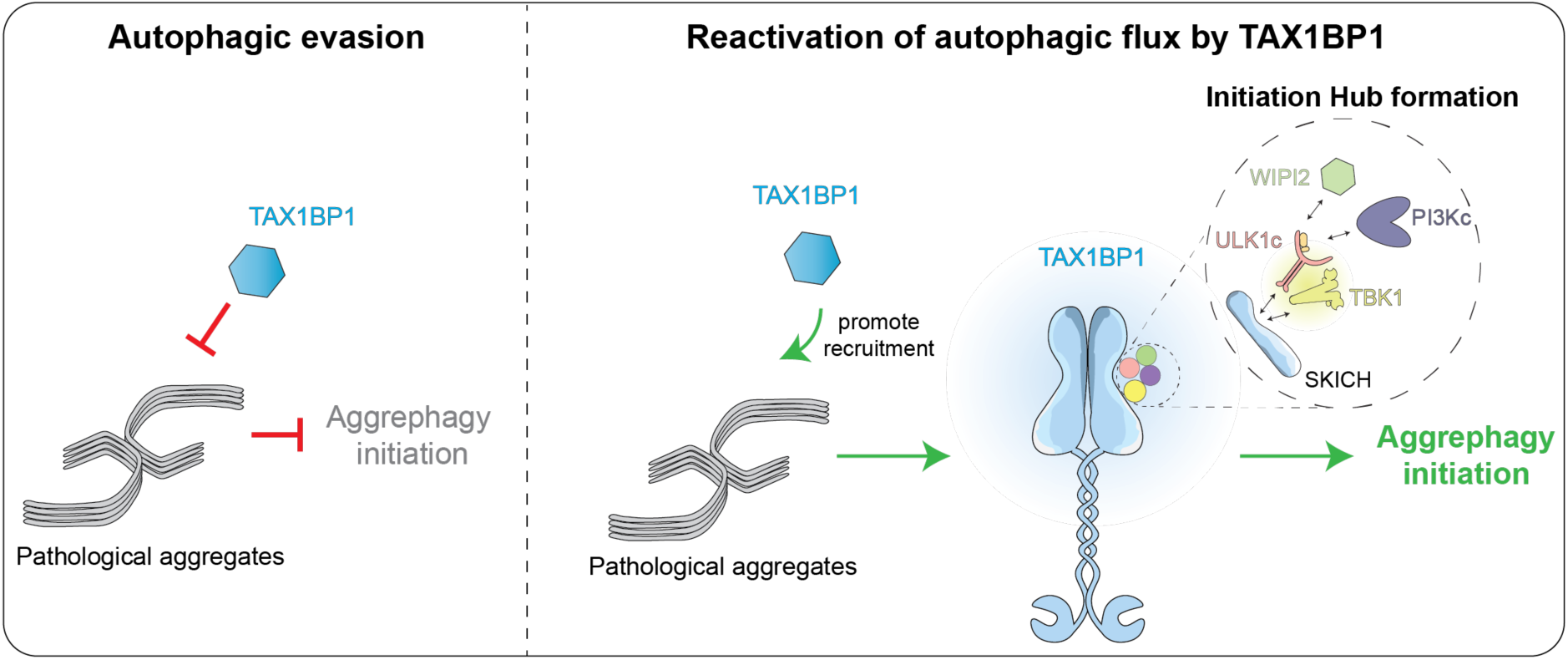
Co-Clustering of TBK1 and ULK1c is essential for aggrephagy initiation. Model of autophagy evasion by pathological protein aggregates and its reactivation through TAX1BP1 recruitment and the subsequent clustering of TBK1 and ULK1 complex.

## Discussion

It is generally assumed that autophagy can degrade solid protein aggregates associated with neurogenerative disease such as those formed by Tau, alpha-synuclein or huntingtin. This assumption originates from the finding that inhibition of autophagy results in the accumulation of protein aggregates in brains and neurons. Moreover, human genetics links loss of function of autophagy proteins with neurodegeneration, while activation of autophagy alleviates aggregate burden ^55–61^. Indeed, the ability of autophagy to sequester large cellular structures in bulk provides a plausible mechanism for its aggregate clearing capacity.

However, it is emerging that solid aggregates are not good substrates for autophagy, as multiple studies found that these aggregates are either poorly or non-productively targeted by the autophagy machinery ^4–8^. It has also been found that aggregates require fragmentation prior to their sequestration by autophagosomes ^62^.

While the Tau aggregates are decorated by p62, they exclude the autophagy initiator TAX1BP1 (Fig. 2), analogous to what was found *in vitro* for Tau fibrils extracted from post-mortem AD brains ^4^. Here, we show that this exclusion is the critical step in autophagic evasion of protein aggregates, because artificially tethering TAX1BP1 to these aggregates restores their degradation. Thus, the recruitment of TAX1BP1 appears to be the breaking point in the activation of autophagy at the aggregates. We have previously shown that the recruitment of TAX1BP1 to ubiquitinated cargoes requires NBR1 as well as ubiquitin binding ^19^. We have further hypothesized that TAX1BP1 is outcompeted for ubiquitin binding by the oligomeric p62:NBR1 heterocomplex, which results in its exclusion from the cargo ^4^. Therefore, enhancing TAX1BP1 recruitment, for example by strengthening the interaction of TAX1BP1 with NBR1 might help to overcome this aggrephagy block.

ULK1 and TBK1 are two major kinases triggering autophagy initiation downstream of TAX1BP1 ^28^. TBK1 is the master regulator of optineurin-driven mitophagy ^29^. The TAX1BP1 related cargo receptor NDP52 can initiate mitophagy through either ULK1 or TBK1 ^29^. By contrast, TAX1BP1-mediated aggrephagy requires the activity of both kinases. This suggests that distinct initiation requirements exist for different substrates. In line with this, the individual recruitment of TBK1 or the ULK1 complex supports mitophagy ^6,63^, but is not sufficient to trigger aggrephagy. The differential requirement for the initiation of autophagic degradation of mitochondria and the Tau aggregates is surprising, because both are large, non-liquid cargoes. However, an important distinction between them may lie in the properties of their surface, as has been put forward previously ^6^. Unlike mitochondria which represent a cargo with a fluid-like membrane surface, allowing dynamic surface interactions and productive autophagy machinery clustering, Tau protein aggregates represent a rigid structure consisting of several polypeptide chains aligned in a higher-order. However, once TAX1BP1 decorates the surface of the Tau aggregates, it potentially allows the formation of dynamic interaction hubs through local clustering of the autophagy machinery on its SKICH domain.

Overall, our study advances the understanding of the molecular mechanism of why pathological aggregates form poor substrates for autophagy. Our finding that TAX1BP1 recruitment is sufficient to drive the delivery of Tau aggregates to lysosomes, demonstrates its critical role in this pathway and underscores the aggregate clearance potential of autophagy in a disease-relevant context. Moreover, the discovery that clustering of TBK1 and the ULK1 complex by TAX1BP1 is the mechanistic trigger for aggrephagy initiation opens new therapeutic avenues for targeting neurodegenerative diseases.

## Acknowledgements

We thank Elias Adriaenssens, Luca Ferrari and Maximilian Schmid for comments on the manuscript. We thank Susanna Tulli for help with deconvolution, and we thank Lea Radzuweit for helping to set up the Tau seeding assays. We also thank Luca Ferrari for the help setting up the Tau fibrillization. We thank the Max Perutz Labs BioOptics, Flow Cytometry, and Mass Spectrometry facilities for technical support. Proteomics analyses were performed by the Mass Spectrometry Facility at Max Perutz Labs using the VBCF instrument pool. We thank the Vienna BioCenter Core Facilities (VBCF) Protech Facility for help with HEK cell expressions. This work was supported by the Austrian Science Fund (FWF Grant-DOI 10.55776/F79 and 10.55776/PAT7165623) and the Vienna Science and Technology Fund (WWTF, grant LS21-015).

## Author contributions

S.M. supervised the overall project. S.M. and B.B. conceived the experiments. B.B. performed the experiments. M.S. assisted with the experiments and provided technical help. S.M. and B.B. wrote the manuscript and participated in all data analysis and interpretation.

## Methods

### Reagents

The following chemicals were used in this study: TBK1 inhibitor GSK8612 (S8872, Selleck Chemicals), ULK1/2 inhibitor (MRT68921, BLDpharm), Vps34-IN1 inhibitor (APE-B6179, ApexBio), A/C hetero-dimerizer (635057, Takara), Bafilomycin A1 (sc-201550, Santa Cruz Biotech), Q-VD-OPh (A1901, ApexBio) and dimethylsulfoxide (DMSO; D2438, Sigma).

### Clonings

The sequences for the insert cDNAs were generated through amplification from existing vectors. The target vectors were opened through restriction enzyme digestion or PCR amplification. All newly generated plasmids were assembled through Gibson cloning using the 2x NEBuilder HiFi DNA assembly enzyme mix (M5520A, New England Biolabs) and subsequently transformed into competent DH5a *E.coli*. The next day single colonies were picked, grown overnight, followed by plasmid isolation using the GeneJet Plasmid Miniprep Kit (K0503, Thermo Fisher). To verify the correct insertion, the plasmids were submitted to Sanger sequencing (MicroSynth AG).

### Cell lines

All cell lines were cultured at 37°C in a humidified 5% CO*_2_*. HeLa (RRID: CVCL_0058) and HEK293T (RRID: CVCL_0063) cells were acquired from the American Type Culture Collection (ATCC). HeLa 5KO (RRID: CVCL_C2VN) and FIP200 KO (CVCL_D1LA) cell lines have been previously generated. HeLa TBK1 KO cell line was obtained by Michael Lazarou (SMcl 134). HeLa and HEK293T cells were grown in Dulbecco Modified Eagle Medium (DMEM, 41966-029, Thermo Fisher) supplemented with 10% (v/v) fetal bovine serum (FBS, Thermo Fisher), 25 mM HEPES (15630080, Thermo Fisher), 1% (v/v) non-essential amino (NEAA, 11140050, Thermo Fisher), and 1% (v/v) penicillin–streptomycin (15140122, Thermo Fisher).

### Generation of stable cell lines

To generate stable cell lines HEK293T were transfected with VSV-G, GAG POL and lentiviral plasmids containing the gene of interest using Lipofectamine 3000 (L3000008, Thermo Fisher). After 48 hours, the viral supernatant was collected and filtered to avoid cross-contaminations. The viral supernatant was added to target cells together with polybrene (8mg/ml) (TR-1003-G, Sigma). After 24 hours, the viral supernatant was removed and replaced with virus-free medium.

The following viral vectors were used in this study: GAG-POL (gift from Versteeg lab; SMc 1768), VSV-G (a gift from Versteeg lab; SMc 1767), pSicoR-Tau.K18-mRuby2 (Addgene_133057), pSicoR-Tau.K18-mClover2 (Addgene_133058), pSicoR-Tau.K18-mKeima-FRB (SMc 2689), pHAGE-FKBP-GFP-ATG13 (SMc 2573), pHAGE-FKBP-GFP-TBK1 (SMc 2100), pHAGE-TAX1BP1-GFP-FKBP (SMc 2836), pHAGE-TAX1BP1(ΔSKICH)-GFP-FKBP (SMc 2906), pHAGE-TAX1BP1(ΔZnF)-GFP-FKBP (SMc 2837), pHAGE-TAX1BP1(ΔFIP200)-GFP-FKBP (SMc 2904), pHAGE-FKBP-BFP-TBK1 (SMc 2944), pHAGE-FKBP-BFP (SMc 2945), pGenLenti-mKeima-p62 (SMc 2775), pGenLenti-mClover-p62 (SMc 2663), pGenLenti-mClover-p62ΔPB1 (SMc 2664), pGenLenti-mClover-p62ΔUBA (SMc 2669)

### Protein production

The SKICH-GST (SMc 1809), SKICHΔFIP200-GST (SMc 1894), GST-TEV-Tau.K18(P301L) (SMc 2541) and WIPI2-mCherry protein (Addgene_223725) were all expressed in bacteria. To this end, the plasmids were transformed into E. coli Rosetta pLysS cells (71403-3, Novagen). For WIPI2-mCherry (Addgene_223725), SKICH-GST (SMc 1809) and SKICHΔFIP200-GST (SMc 1894), the transformed bacteria were grown in 2x tryptone yeast extract (TY) medium at 37 °C until reaching an optical density at 600 nm (OD600) of 0.4 and then continued at 18 °C. When the bacterial culture reached an OD600 of 0.8, 100 µM isopropyl β-D-1-thiogalactopyranoside (367-93-1, Gerbu) was added to induce protein expression, and the cultures were incubated for 18 hours at 18°C. The bacteria were harvested by centrifugation, and the pellets were frozen in liquid nitrogen and stored at -70°C. For GST-TEV-Tau.K18(P301L) (SMc 2541), the transformed bacteria were grown in 2x tryptone yeast extract (TY) medium at 37 °C until reaching an optical density at 600 nm (OD600) of 0.6. 10 mM Betaine and 200 µM isopropyl β-D-1-thiogalactopyranoside was added to the cultures for effective protein production. The cultures were incubated for 3 hours at 30°C and subsequently harvested by centrifugation. The pellets were frozen in liquid nitrogen and stored at -70°C

The PI3Kc-mCherry (Addgene_187936), FIP200-GFP (Addgene_187832) and GFP-TBK1 (Addgene_187830) proteins were expressed in Sf9 insect (12659017, RRID: CVCL_0549, ThermoFisher). To generate the bacmid DNA for insect cell expressions, we transfected our plasmids into DH10EMBacY cells using electroporation. The successful insertion of our gene of interest was validated by PCR and the bacmid DNA was subsequently isolated. 5 µg of bacmid DNA was transfected into Sf9 cells using FuGene HD (E2311, Promega) and the expression of YFP was monitored over several days. After five to seven days the supernatant was collected (V0) and used to infect 30 ml of Sf9 culture at one million cells per ml. After five to seven days and a significant drop in cell viability and a confirmed YFP signal, the V1 viral supernatant was harvested and filtered. 1 ml of V1 was used to infect 1 liter of Sf9 cells. The cells were harvested once the cell viability dropped to around 90% (typically 3-5 days after infection). The cells were harvested by centrifugation, and the pellets were frozen in liquid nitrogen and stored at -80°C.

The NAP1-mCherry (Addgene_198036) and ULK1c-GFP (Addgene_171410, Addgene_171413, Addgene_189590) proteins were expressed by the VBCF Protech facility (https://www.viennabiocenter.org/vbcf/protein-technologies/). The proteins were expressed in FreeStyle HEK293F cells, grown at 37 °C in FreeStyle 293 expression medium (Thermo Fisher). The day before transfection, cells were seeded at a density of 0.7 × 10^6^ cells ml^−1^. On the day of transfection, a 400-ml culture was transfected with 400 μg of the MAXI-prep DNA, diluted in 13 ml of Opti-MEM I reduced serum medium (Thermo Fisher) and 800 μg of polyethylenimine (PEI 25K, Polysciences), also diluted in 13 ml of Opti-MEM media. One day post transfection, the culture was supplemented with 100 ml of EXCELL 293 serum-free medium (Sigma-Aldrich). Another 24 h later, cells were collected by centrifugation at 270g for 20 min. The pellet was washed with PBS to remove medium, then flash-frozen in liquid nitrogen. Pellets were stored at −70 °C.

### Protein purification

To purify SKICH-GST (SMc 1809) and the SKICHΔFIP00-GST (SMc 1894), the pellets were resuspended in 30 ml of lysis buffer (300 mM NaCl, 50 mM HEPES pH 7.5, 1 mM DTT, 2 mM MgCl2, cOmplete EDTA-free protease inhibitor (Roche), DNase (Sigma)) and lysed through sonication. Afterwards the lysate was centrifuged for 45 minutes with 18 000 rpm at 4°C to separate soluble from insoluble fractions. The supernatant was mixed with 3 ml of Glutathione Sepharose 4B beads (17075605, Cytiva) which were previously washed and equilibrated. The lysate and the beads were incubated for 90 minutes rolling at 4°C. Then the beads were spun down, the supernatant discarded and the beads washed 5 times with washing buffer I (50 mM HEPES, pH 7.5, 300 mM NaCl, 1 mM DTT), 1 x with washing buffer II (50 mM HEPES, pH 7.5, 700 mM NaCl, 1 mM DTT) and again washed twice with washing buffer I. The protein was eluted by incubating the beads with elution buffer (Wash Buffer I + 50 mM glutathione (Sigma) - pH adjusted to 7.5) overnight at 4°C. The next day, the eluate was collected by centrifugation and subsequently filtered to remove any beads from the solution. The eluate was concentrated to 500 µl using concentrators (Merck Millipore). Then the sample was further purified by size-exclusion chromatography in SEC buffer (25 mM HEPES pH 7.5, 150 mM NaCl, 1 mM DTT). The fractions were validated by SDS-PAGE, pooled and snap-frozen by liquid nitrogen. All proteins were stored at -70°C.

To purify GST-TEV-Tau.K18(P301L) (SMc 2541), the pellets were resuspended in lysis buffer (300 mM NaCl, 50 mM HEPES pH 7.5, 1 mM DTT, 2 mM MgCl2, cOmplete EDTA-free protease inhibitor (Roche), DNase (Sigma)) and lysed through sonication. The lysate was cleared by a centrifugation step at 18 000 rpm at 4°C for 30 minutes. The supernatant was incubated with 5 ml of equilibrated GSH beads (17075605, Cytiva) for 2 hours at 4°C. The beads were washed 3x with washing buffer I (50 mM HEPES, pH 7.5, 45 mM NaCl, 1 mM DTT) and twice with SP Buffer (25 mM HEPES, pH 7.5, 300 mM NaCl, 1 mM DTT). For protein elution, the beads were incubated with 100 µl of TEV protease (2 mg/ml) for 2 hours at 16°C. The eluate was collected, filtered and subsequently applied to an HiTrap SP HP column (17115101, Cytiva) and eluted with a linear gradient (45 mM NaCl – 500 mM NaCl). The eluate was collected and further purified by size exclusion chromatography in SEC buffer (25 mM HEPES pH 7.5, 150 mM NaCl, 1 mM DTT). Individual fractions were validated by SDS-PAGE. Fractions containing the protein of interest were pooled, snap frozen in liquid nitrogen and stored until further use at -70°C.

A protocol for the purification of mCherry tagged WIPI2 (Addgene_223725; https://doi.org/10.17504/protocols.io.4r3l2qqyql1y/v1), mCherry tagged PI3K-C1 (Addgene_187936; dx.doi.org/10.17504/protocols.io.8epv59mz4g1b/v1), GFP tagged TBK1 (Addgene_187830; https://doi.org/10.17504/protocols.io.81wgb6wy1lpk/v1), mCherry tagged NAP1 (Addgene_198036; https://doi.org/10.17504/protocols.io.5jyl8jw6dg2w/v1), GFP tagged FIP200 (Addgene_187832; https://doi.org/10.17504/protocols.io.dm6gpbkq5lzp/v1) and GFP tagged ULK1 complex (Addgene_171410, Addgene_171413, Addgene_189590; https://doi.org/10.17504/protocols.io.bvn2n5ge) have been previously published.

### *In vitro* Tau fibril formation

To induce fibrilization of recombinant Tau, the protein was mixed at a final concentration of 20 µM together with 5 µM Heparin (SantaCruz Biotech) in SEC Buffer (25 mM HEPES pH 7.5 150 mM NaCl, 10 mM DTT, cOmplete EDTA-free protease inhibitor (Roche)) and agitated overnight at 37°C. The next day, the protein solution was sonicated in water 5x with 1 min pulses, snap frozen in liquid nitrogen and stored at -80°C.

### Tau seeding assay

Cells were seeded on 6 well plate a day prior to seeding with the fibrils. For Tau seeding, Tau fibrils were transfected into the cells using Lipofectamine 3000 at a final concentration of 20 nM of fibrils and 4 µl of Lipofectamine per well.

### Pelleting assay

Tau seeding was performed as described above. Cells were harvested by trypsinization and lysed in lysis buffer (150mM NaCl, 50 mM Tris-HCl pH 7.4, 1 mM EGTA, 1 mM EDTA, 1% Triton X-100, 0.27 M sucrose, 1 mM DTT, cOmplete EDTA-free protease inhibitor (11836170001, Roche)). After a 30 min incubation on ice, the lysates were subsequently cleared by two centrifugation steps with 500 x g (5 min) and 1000 x g (5 min) at 4°C. After taking an Input sample, the cleared lysates were submitted to ultracentrifugation for 1 hour with 100 000 x g at 4°C. The supernatant was collected, and the pellet was washed 1x with PBS and spun for 30 min with 100 000 x g. The supernatant was discarded, and the pellet was resuspended with the volume and lysis buffer as the original lysate, from which an input sample was taken. The samples were analyzed through SDS-PAGE and western blotting as described below.

### Rapalog-induced dimerization experiments

For all rapalog-induced tethering experiments, the cells were first seeded with the Tau fibrils to induce the aggregation of the over-expressed Tau in the cells. The next day, the cells were treated with 500 nM Rapalog (Takara) for either 24 or 48 hours. For experiments, that include the HeLa 5KO and FIP200 KO cell lines, 10 μM Q-VD-OPh (ApexBio) was added to suppress apoptosis.

### Flow Cytometry

Tau aggregation and Rapalog induced tethering was performed as described above. Cells were harvested by trypsinization and then filtered through 35 µm cell-strainer caps (352235, Falcon). The samples were analyzed by an LSR Fortessa Cell Analyzer (BD Biosciences). Lysosomal Tau.K18-mKeima signal was measured using dual excitation ratiometric pH measurements at 405 nm (pH 7) and 561 nm (pH 4) lasers with 710/50 nm and 610/20 nm detection filters. For the tethering experiments, GFP was used as an additional channel. For the co-tethering of ATG13 and TBK1, GFP and BFP were used as additional channels. For each experiment 100 000 Tau.K18-mKeima positive cells were analyzed. The gates for the aggrephagy positive population were drawn based on the control sample that was treated with Tau aggregates, Rapalog and Bafilomycin. All Flow Cytometry experiments were analyzed using FlowJo (SCR_008520).

### SDS-Page and western blot analysis

Except for the pelleting assays (see above), cells were harvested by trypsinization and lysed in RIPA buffer (50 mM Tris-HCl pH 8.0, 150 mM NaCl, 0.5% sodium deoxycholate, 0.1% SDS, 1% NP-40) supplemented by cOmplete EDTA-free protease inhibitors (11836170001, Roche) and phosphatase inhibitors (Phospho-STOP; 4906837001, Roche). After 30 minutes on ice, the lysates were cleared by a centrifugation step at 13 000 rpm for 10 min (4°C). Protein concentrations were measured by Bradford (23246, ThermoFisher). Equal protein amounts were loaded on 4%-12% SDS-PAGE gels (ThermoFisher) together with PageRuler Prestained protein marker (Thermo Fisher), except for the LC3B blots, which were analyzed with 16% Tris-glycine gels (Thermo Fisher). The proteins were subsequently transferred onto a nitrocellulose membrane (10600001, Cytiva) for 1 h at 4 °C using the Mini Trans-Blot Cell (Bio-Rad). After the transfer the membranes were blocked with 3% milk powder dissolved in TBS-T (0.1% Tween 20 in TBS) for 1 hour at RT. The membranes were subsequently incubated with the primary antibodies (diluted in blocking solution) overnight at 4°C. The next day, the blots were washed 3x witch TBS-T and then incubated with secondary antibodies fused to HRP (diluted 1:10 000 in blocking solution) for 1 hour at RT. After 3 TBS-T washings steps, the blots were developed using Super Signal West femto chemiluminescence substrate (34096, Thermo Fisher) and imaged with a ChemiDoc MP Imaging system (Bio-Rad).

The following primary antibodies and dilution were used in this study: rabbit anti-Tau (1:2500; MA5-42451, Invitrogen), mouse anti-p62 (1:1000; #610832, Clone 3/p62 lck ligand, BD Bioscience), mouse anti-NBR1 (1:1000; H00004077-M01, Clone 6B11, Abnova), rabbit anti-TAX1BP1 (1:1000; #5105, clone D1D5, CST), rabbit anti-FIP200 (1:1000; #12436, clone D10D11, CST), rabbit anti-TBK1 (1:1000; #3013, CST), rabbit anti-TBK1 pS172 (1:1000; #5483, clone D52C2, CST), rabbit anti-ATG13 (1:1000; #13468, clone E1Y9V, CST), rabbit anti-ULK1 (1:1000; #8054, clone D8H5, CST), mouse anti-LC3B (1:1000; #3868S, clone D11, CST), rabbit ATG13 p-S355 (1:1000; #46329, CST), rabbit anti ULK1 p-S317 (1:1000; #37762, CST), rabbit anti-ULK1 p-S757 (1:1000; #14202, CST), rabbit anti-ATG9 (1:1000; #13509, clone D4O9D, CST), rabbit anti-ATG16L1 (1:1000; #8089, clone D6D5, CST), rabbit anti-ATG14 (1:1000; #5504, CST), mouse anti-GAPDH (1:20000; #G8795, clone 71.1, Sigma) The secondary antibodies used in this study were horseradish peroxidase (HRP)-conjugated polyclonal goat anti-mouse (1:10000, #115-035-003, Jackson ImmunoResearch Labs) and HRP-conjugated polyclonal goat anti-rabbit (1:10000; #111-035-003, Jackson ImmunoResearch Labs).

For quantification, a rectangle was drawn around the corresponding protein band in ImageJ and the intensity was plotted using the “Plot Lanes” function. The measured values were adjusted by using GAPDH as a loading control. For experiments analyzing the phosphorylation level of proteins, signals of the phospho-specific antibodies were adjusted to the signal of the total protein.

### Immunofluorescence and confocal microscopy

Cells were seeded on glass cover slips (0117520, Marienfeld Superior) and the aggregation of Tau and the rapalog-induced tethering were performed as described above. Cells were washed 3x with PBS and subsequently fixed with 4% Paraformaldehyde (in PBS) for 20 minutes - except for stainings for LC3B, for which the coverslips were fixed in ice-cold Methanol for 20 minutes on ice. After three washing steps with PBS, the cells were treated with 0.1% Triton (in PBS) for 5 minutes to permeabilize the cell membranes - except for stainings for FIP200. After two PBS washing steps, the coverslips were blocked with 1% BSA (in PBS) for 1 hour. Then the coverslips were incubated for 1 hour with the primary antibody (diluted in blocking solution). After three PBS washing steps, the coverslips were incubated with secondary antibodies (diluted in blocking solution). The coverslips were incubated for 1 hour with the secondary antibody and subsequently mounted on glass sides (AA00000112E01MNZ10, epredia) using DAPI-Fluoromont-GTM (0100-200, Southern Biotech).

For FIP200 staining, the coverslips were permeabilized after fixation with 0.25% Triton (in PBS) for 15 minutes. Subsequently, the coverslips were incubated with the primary antibody (in blocking solution) for 1 hour at 37°C, washed 3x with PBS and then incubated with the secondary antibody again for 1 hour at 37°C. Afterwards, the coverslips were treated as described above.

Deconvolution was performed for Figure 3F with confocal settings, a total of 40 iterations, and the signal-to-noise ratio parameter was set to 20. Output was generated as 32-bit ICS2 files for further processing with Fiji ImageJ.

The following primary antibodies and their dilutions were used in this study: rabbit anti-Tau (1:500; MA5-4245, Invitrogen), mouse anti-p62 (1:100; #610832, Clone 3/p62 lck ligand, BD Bioscience), rabbit anti-TAX1BP1 (1:100; #5105, clone D1D5, CST), rabbit anti-p62 p-S403 (1:100; #39786, clone D8D67, CST), rabbit anti-FIP200 (1:200; #12436, clone D10D11, CST), rabbit anti-TBK1 p-S172 (1:100; #5483, clone D52C2, CST), rabbit anti-ATG13 (1:100; #13468, clone E1Y9V, CST), rabbit anti-ULK1 (1:100; #8054, clone D8H5, CST), mouse anti-LC3B (1:100; 0260-100/LC3-2G6, NanoTools), mouse anti-ubiquitin (UBCJ2 (1:500; ENZ-ABS840, Enzo Life Sciences).

The following secondary antibodies were used in this study: AlexaFluor-488 goat anti-mouse immunoglobulin G (IgG) (H + L) (1:1000, A-11001, Invitrogen), AlexaFluor-488 goat anti-rabbit immunoglobulin G (IgG) (H + L) (1:1000, A-11008, Invitrogen), AlexaFluor-647 goat anti-mouse immunoglobulin G (IgG) (H + L) (1:1000, A-21235, Invitrogen), AlexaFluor-647 goat anti-rabbit immunoglobulin G (IgG) (H + L) (1:1000, A-21244, Invitrogen).

Imaging was performed with a Zeiss LSM 700 confocal microscope equipped with a Plan-Apochromat 63×/1.4 Oil DIC objective. Multiple areas within samples were imaged to obtain comparable numbers of cells (50–150 cells). To avoid cross-contamination between fluorochromes, each channel was imaged sequentially using the multitrack recording module before merging. Images from fluorescence and confocal acquisitions were processed and analyzed with ImageJ software.

To quantify the number of aggregates, a previously described macro was used. In short, binary pictures were generated by thresholding in ImageJ. The „Analyze puncta“ function was used to count the number of puncta. The threshold was adjusted manually for each and kept consistent throughout an experiment. Additionally, a suitable pixel units cut-off was used and kept consistent throughout an experiment. To analyze the degree of colocalization between different channels, the Pearson’s correlation coefficient (PCC) was measured using a plug-in called JACoP ^64^. A PCC value of 1 indicates complete colocalization and a PCC value of near 0 means random distribution. To calculate the PCC, individual cells containing Tau aggregates were marked and added to the ROI manager in Fiji to exclude cells without aggregates from the analysis.

### Microscopy-based protein-protein interaction assay

Glutathione Sepharose 4B beads (17075601, Cytiva) were first washed with water and then equilibrated in SEC Buffer (150 mM NaCl, 25 mM HEPES pH 7.5 1mM DTT). The beads were subsequently incubated with 5 µM bait protein for 1 hour at 4°C. Afterwards the beads were washed twice with SEC buffer and then mixed with a prey protein solution (each prey protein at 500 nM) in a 384 well plate. After 30 minutes of incubation to reach an equilibrium, the beads were imaged with a Zeiss LSM 700 confocal microscope equipped with Plan-Apochromat 20X/0.8 WD 0.55 mm objective.

### Statistics

All experiments have been performed at least three times. Statistical analyses were performed using the statistical tests stated in each figure legend. Data are presented as mean +/-standard deviation for immunoblotting or flow cytometry-based quantification and as violin plots for microscopy-based quantifications. For microscopy-based quantification individual data points represents either individual cells (Pearson correlation coefficient) or the average of a single image (quantification of number of aggregates). Immunoblotting and flow cytometry-based quantification, each data points represent the corresponding measurements of the replicate. Microscopy images and immunoblots were quantified using ImageJ. For flow cytometry quantification we used FlowJo. All graphs were generated by GraphPad Prism version 9.5.1. All statistical tests have been performed in GraphPad Prism. Depending on the number of samples, we performed a t-test, one- or two-way ANOVA test with appropriate multiple comparison tests. All P values below 0.05 were considered significant, with *P ≤ 0.05, **P ≤ 0.01, ***P ≤ 0.001 and ****P ≤ 0.0001. For Fig. 4h-j, random positions based on DAPI staining were imaged to avoid biased imaging. For all other experiments, no blinding or randomization was performed.

## Supplemental Figures

**Figure S1:**
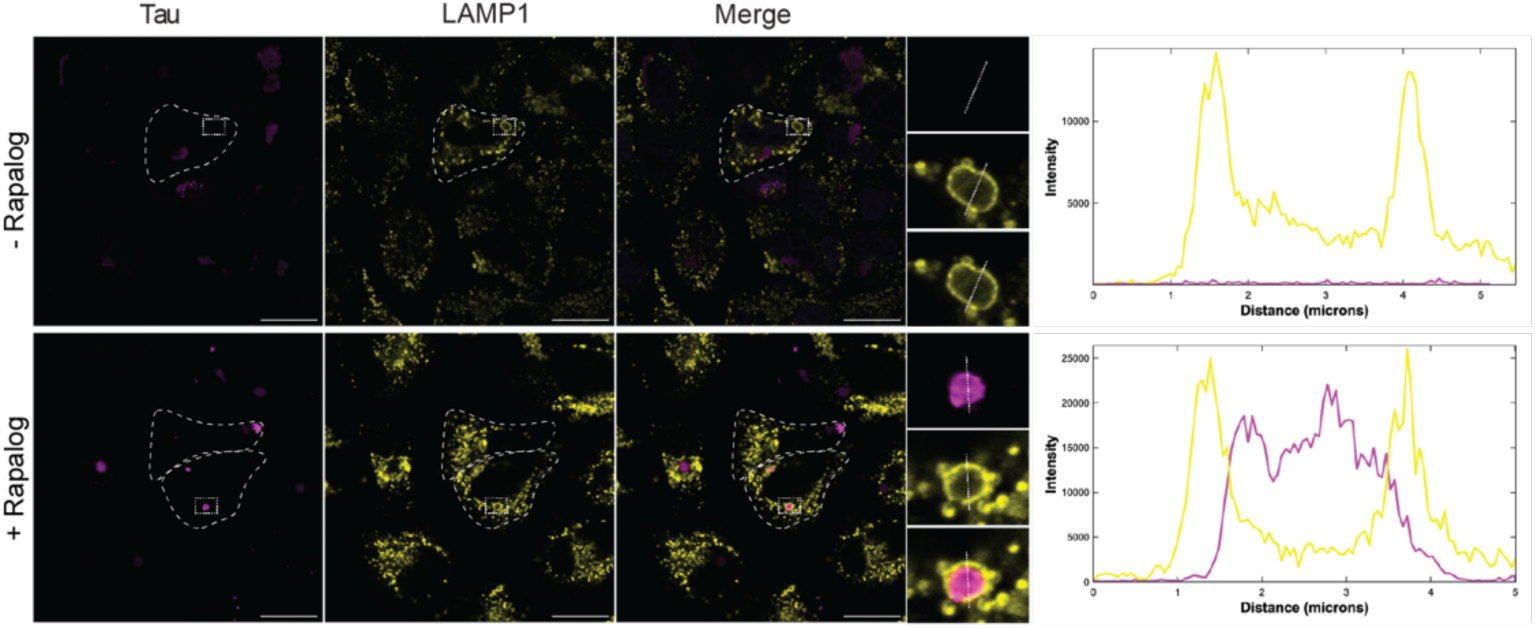
Tau is delivered into lysosomes upon TAX1BP1 tethering. Immunofluorescence images of Tau and the lysosomal marker LAMP1 with and without rapalog treatment. Line diagram of fluorescence signal of LAMP1 and Tau along the indicated line to the right side of the individual images.

**Figure S2:**
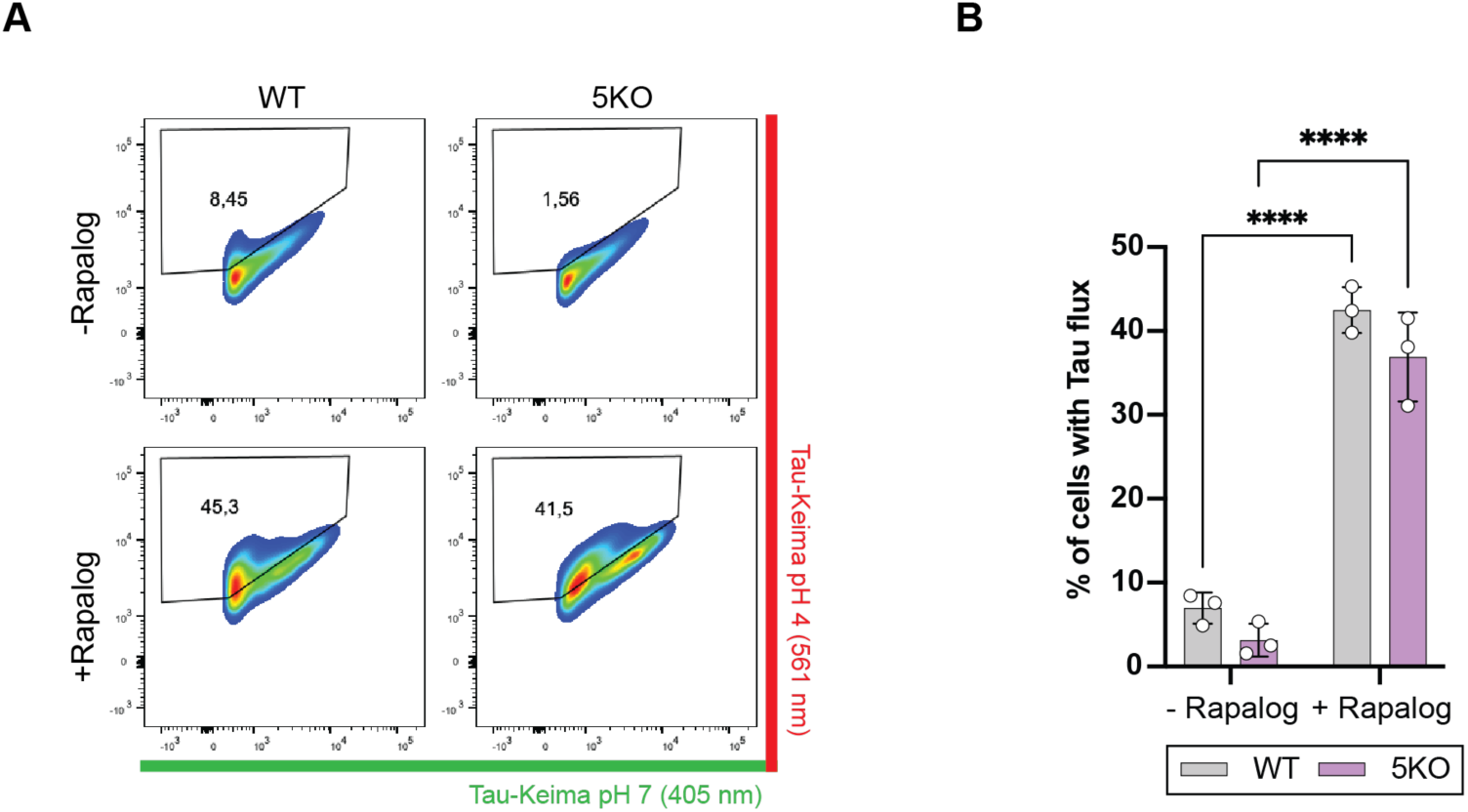
TAX1BP1 is sufficient to induce autophagic Tau aggregate flux. **A)** Representative fluorescence-activated cell sorting (FACS) plots of Tau flux measurements in WT and 5KO cells upon TAX1BP1 tethering. **B)** Comparison of the Tau flux in WT and 5KO cells upon TAX1BP1 tethering. Data are presented as mean +/- s.d. (n = 3 biologically independent experiments). Two-way ANOVA with Sidak’s multiple comparisons test. ****P < 0.0001. ns, not significant.

**Figure S3:**
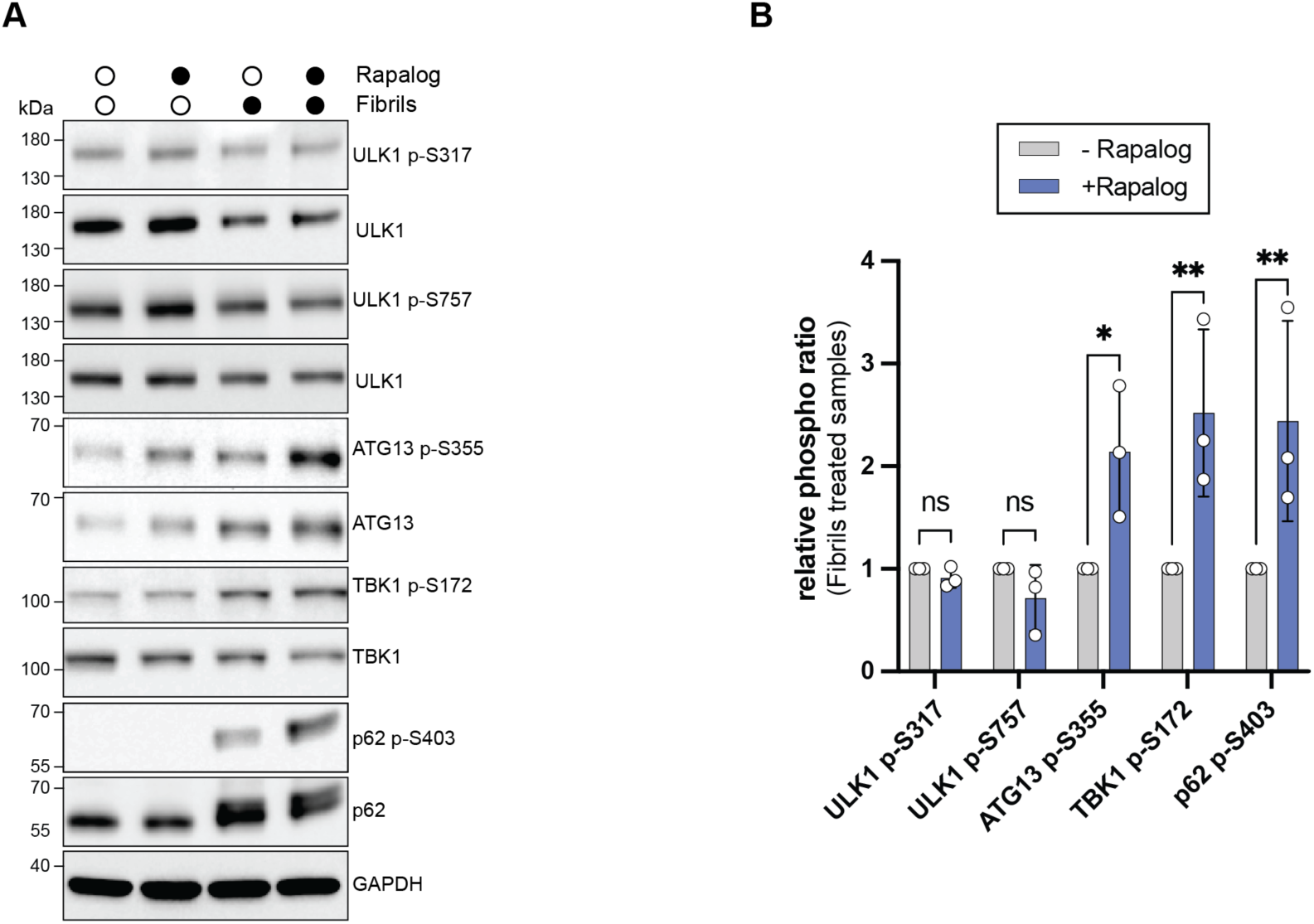
TAX1BP1 activates the ULK1 complex and TBK1. **A)** Immunoblotting of phospho-ULK1, ATG13, TBK1 and p62 upon Tau aggregation and TAX1BP1 tethering. **B)** Quantification of relative increase of phosphorylation levels upon TAX1BP1 tethering. Phosphorylation signal was normalized to the total protein. To account for inter-experimental variability, values are expressed as a relative to the -Rapalog samples within each biological replicate. The data are presented as mean +/- s.d. (n = 3 biologically independent experiments). Two-way ANOVA with Sidak’s multiple comparisons test. *P < 0.05, **P < 0.005. ns, not significant.

